# The impact of late Pleistocene mammal extinctions on pathogen richness in extant hosts

**DOI:** 10.1101/2023.07.18.549351

**Authors:** Tomos O. Prys-Jones, Andrew J. Abraham, Joseph R. Mihaljevic, Kris A. Murray, Christopher E. Doughty

## Abstract

Many species of large mammals were driven to extinction during the late Pleistocene and early Holocene (approx. 10,000 – 50,000 years ago), with cascading effects on the physical structure of ecosystems and the dispersal of seeds, nutrients, and microbes. However, it remains uncertain whether the parasites associated with these extinct hosts also disappeared or persisted in surviving (extant) mammals. We hypothesize that if some parasites endured, extant mammals sharing their ranges with phylogenetically similar extinct mammals would have a greater pathogen richness than expected based on current levels of host diversity. We find that the inclusion of variables related to these extinctions account for an additional 5% of deviance when modelling per-host viral and bacterial richness, compared to models run without these variables. Partial dependence plots show a positive correlation between the number of extinct mammals lost and per-host viral and bacterial richness (p < 0.001 and p = 0.03, respectively). Additionally, decreasing phylogenetic distance between the extinct and extant species is associated with an increasing viral richness (p < 0.001). We discuss four mechanisms that may be driving these patterns and highlight future research to distinguish between them. Next, we use the models and IUCN range maps to identify geographic regions where viral and bacterial richness differs due to the inclusion of extinction variables. Notably, the richness of both pathogen types is increased in South America (viruses: +6.8%; bacteria: +3.1%) and decreased in Africa (viruses: −2.6%; bacteria: −13.6%), two continents known to have high and low levels of historical mammal extinctions, respectively. Viral richness is also elevated in North America (+8.6%), Europe (+5.1%), Oceania (+3.3%), and Asia (+2.3%). These results support the inclusion of extinction variables in future models of pathogen richness and may allow for improved targeting of future surveillance efforts.

## INTRODUCTION

The recent SARS-CoV-2 pandemic highlighted a lack of awareness of the pathogens circulating in wildlife populations (Guo et al., 2020; Hraib et al., 2022; Kaler et al., 2022) and emphasized a need for greater surveillance. Estimates suggest that 16.9% of mammal species (Burgin et al., 2018) and only 2.1% of their viruses (Carlson et al., 2019) are contained within datasets that record the associations between host and parasite species (Gibb et al., 2021).

Many repositories of host-parasite associations currently exist, collated using a variety of methodologies. Notably, Shaw et al. (2020) performed a systematic literature review to compile a database of associations between 2,656 vertebrate host species and 2,595 viral or bacterial pathogens. A similar process was used in generating the HP3 viral dataset used by Olival et al. (2017). The studies included in these manual literature searches used a variety of methods to detect the pathogen in the host species, including polymerase chain reaction (PCR) and serological methods (viral or serum neutralization tests, enzyme linked immunoassays, antigen detection assays). Recently, both datasets have been incorporated into the large CLOVER database (Gibb et al., 2021).

These datasets allow for effective targeting of surveillance to specific taxa or regions of the world that have a greater number of parasites. Ideally, all parasites would be characterized in all host species, but this is not possible at present due to a lack of resources. Instead, studies use the subset of host-parasite associations available in repositories and identify host traits that correlate with their parasite richness. (Olival et al., 2017) and (Shaw et al., 2020) found that the biological variables associated with the per-host richness of bacteria or viruses included body mass, host range, and mammal sympatry. In addition to variables related to host ecology, research effort is consistently identified as being positively correlated with pathogen richness. (Olival et al., 2017) then ran their fitted models with an elevated level of research effort to understand the expected viral richness one would predict given further screening of the mammal species included in the study. Using the range maps of these host species, the authors were able to identify geographic regions with elevated viral richness, and therefore where surveillance efforts would be most effective at finding novel viruses.

In addition to the variables used by Olival et al. (2017), pathogen richness is also associated with land use and climate changes. Gibb et al. (2020) and Albery et al. (2022) showed that urban adapted mammals and those with faster life history traits tend to have a greater pathogen richness. These life history traits include gestation length, litter size, neonate body mass, interbirth interval, weaning age, and sexual maturity age (Albery et al., 2022; Plourde et al., 2017). It is generally accepted that a faster life history is associated with greater investment in reproduction over survival and as a result a lower investment in adaptive immunity (Lee et al., 2008; Previtali et al., 2012). Consequently, mammals with a faster life-history tend to host a greater richness of pathogens (Albery et al., 2022). Alterations to viral sharing between hosts are also projected with anthropogenically-driven climate change, as range shifts in mammalian species create new host assemblages and opportunities for pathogen transmission (Carlson et al., 2022).

While current climate and land use change are the focus of recent studies, humans have been modifying ecosystems for thousands of years. Since the Late Pleistocene (130,000 YBP) and the global spread of anatomically modern humans from Africa, anthropogenic pressures have caused widespread mammal extinctions, particularly amongst the larger species, otherwise known as megafauna (Dirzo et al., 2014; Malhi et al., 2016; Sandom et al., 2014; Smith et al., 2018; Stuart, 2015). At least 154 megafauna species (≥ 44 kg body mass) were lost between 132,000 and 1000 years before present (YBP). These taxa are often considered ecosystem engineers (Enquist et al., 2020) and their removal likely disrupted many features of the ecosystem, including its physical structure (Malhi et al., 2016; Zimov et al., 2015), nutrient transport (Doughty et al., 2013; Wolf et al., 2013), food webs (Fricke et al., 2022), and seed (Bunney et al., 2017; Doughty et al., 2016; Guimarães et al., 2008; Pires et al., 2017), microbe and parasite dispersal (Doughty et al. 2020). Larger species also tend to carry a greater diversity of symbiotic gut microbes (Sherrill-Mix et al., 2018) and parasites (Esser et al., 2016; Kamiya et al., 2014).

Accordingly, mammal biodiversity in the present day is not representative of that 10,000 YBP. Stuart (2015) highlights the geographic heterogeneity of the extinctions, with greater numbers of megafauna lost from continents outside of Africa, and particularly in the Americas (Figure S1). While pathogen richness in the present day is correlated positively with the number of mammal and avian species (Dunn et al., 2010), it is not clear whether a tighter correlation would be achieved by also including extinct hosts.

Historical mammalian extinctions are increasingly being recognized as affecting the distributions of individual parasites, but to date have not been included collectively in global models of parasite richness. For example, Farrell et al. (2021) highlighted that a parsimonious explanation for the presence of the elephant tapeworm (*Anoplocephala manubriata*) in both Asian and African elephants (*Elephas maximus* and *Loxodonta africana,* respectively) despite no geographic overlap, is through a consideration of proboscidean extinctions. The estimated distribution for the extinct woolly mammoth (*Mammuthus primigenius*) would have bridged the elephantid populations, potentially allowing the tapeworm to parasitize both.

Parasites predominantly infecting a single extinct Late Pleistocene species would likely have faced co-extinction with the loss of this host. Galetti et al. (2018) use a linear relationship of the number of helminths per host (Poulin & Morand, 2000) to estimate that the loss of 177 mammalian megafauna species during the late Pleistocene could have led to the co-extinction of at least 444 helminth species. However, a portion of the parasites on soon-to-be extinct Late Pleistocene species may have survived through host-switching (Hoberg & Brooks, 2008). Bush and Kennedy (1994) refer to this phenomenon as ‘host capture’, defining it as the acquisition of a host or host group not normally associated with the parasite. The process of capturing a new host is common to both macroparasites and microparasites (Hoberg & Brooks, 2008; Longdon et al., 2014; Perlman & Jaenike, 2003). Generally, the probability of host capture is higher when the phylogenetic distance between the new and original host is low (Albery et al., 2020; Gilbert & Webb, 2007; Streicker et al., 2010), as the internal physiology and ecological niches have less time to diverge (Woolhouse et al., 2012; Woolhouse et al., 2005). However, some have speculated that host capture was prevented during the Late Pleistocene due to the scale of the extinctions and the loss of entire mammal clades, removing all the most suitable potential hosts that a pathogen might use (Galetti et al., 2018).

The extinctions may have also threatened generalist parasites that infected a broader suite of Late Pleistocene species. Generalists unable to sustain a net reproductive rate greater (NRR) than 1 in the post-extinction species would have been lost (Farrell et al., 2015). The extinctions may also have facilitated transmission of ‘apparent specialist’ parasites into new hosts as there would have been many cascading effects in the ecosystem (mentioned above) following the loss of mammal species, allowing novel interactions of the surviving hosts (Young et al., 2014).

For the pathogens of terrestrial mammals that were able to reproduce in new or existing hosts following the Late Pleistocene extinctions, the average dispersal capacity would likely have reduced, as the largest, furthest ranging species were lost. Doughty et al. (2020) estimate a global 7-fold reduction in fecal microbe and ectoparasite dispersal by terrestrial mammals following the extinctions. The authors also found that these estimates of reduced dispersal improved the performance of statistical models in predicting the locations of zoonotic emergent infectious disease events (EIDs) recorded in human populations within the last 80 years.

Based on the logic of host-switching described above, we expect elevated parasite richness in areas that had higher host richness prior to their extinction. To test this hypothesis, we first calculated the number of extinct mammals that were once sympatric with surviving mammal species and calculated their phylogenetic distances from one another. Second, we used statistical models to determine whether our extinction metrics correlated with pathogen richness in current hosts. To our knowledge, this is the first study that evaluates pathogen richness in the context of late Pleistocene extinctions, evaluating multiple hosts at a global scale. Third, we predicted pathogen richness in surviving hosts with research effort set to the highest recorded value in our dataset. Fourth, we generated maps to identify regions of the world where pathogen richness has changed from predictions of prior studies due to the inclusion of the extinction variables. Such maps may be important in identifying geographic regions with more pathogens than expected, guiding future surveillance efforts.

## METHODS

We used generalized additive models (GAMs) to identify variables associated with the richness of viruses and bacteria in extant hosts. The GAMs were previously published by Olival et al. (2017) and subsequently used by Shaw et al. (2020). However, here we include novel metrics associated with historical mammal losses recorded since the Late Pleistocene, which were generated using the Phylacine repository (version 1.2.1) (Faurby et al., 2018). All supporting data, original scripts and scripts modified from those published by Shaw et al. (2020) and Olival et al. (2017) can be found at Zenodo (DOI: 10.5281/zenodo.8066191).

### HOST SPECIES AND PATHOGEN RICHNESS

The extant hosts and trait data used in this study were initially compiled by Olival et al. (2017) and subsequently used by Shaw et al. (2020). All the species were either terrestrial or volant wild mammals. Data on parasite richness in extant hosts came from CLOVER (version 1.0), the largest repository of host-pathogen associations available (Gibb et al., 2021), a combination of records from Shaw et al. (2020), the EcoHealth Alliance’s HP3 (Olival et al., 2017), Global Mammal Parasite Database version 2.0 (GMPD2) (Stephens et al., 2017) and EID2 (Wardeh et al., 2015). The iteration of CLOVER used here contains 25,195 unique host-pathogen associations between 3,724 unique host species and 6,717 pathogens. In this study we selected to use data on viruses and bacteria/rickettsia (hereafter bacteria). The four datasets comprising CLOVER often capture the same host-parasite associations. Therefore, after filtering for the correct pathogen types, the duplicate host-pathogen associations were also removed. This resulted in viral and bacterial richness data for 511 and 241 host species (from 2840 and 1542 unique host parasite interactions), respectively.

Recently, a study of the CLOVER repository found that the four data sources accounted for 4.7% of the variation in host species’ viral diversity (Gibb et al., 2021), leading the authors to err caution when comparing results from studies of different datasets. They state, ‘*when studies report different findings based on slight variation around a significance threshold, readers should therefore wonder whether subtle differences in the underlying datasets might account for such variation*’. One source of variation between the four datasets is the methodologies used in their collation. While datasets from Shaw et al. (2020), HP3 and GMPD2 were collected from manual literature searches, EID2 was generated from the automated scraping of data from two web sources, PubMed (titles and abstracts) and the NCBI Nucleotide Sequence database. To address the potential that the data repository would affect our results, GAMs were fitted to total per-host parasite richness data from CLOVER, as well as a subset of the data from one of the constituent datasets collected using a manual literature search. Data from Shaw et al. (2020) was used to calculate the per-host viral and bacterial richness, resulting in viral and bacterial richness data for 435 and 188 hosts (from 1655 and 659 unique host parasite interactions), respectively.

### EXTINCTION METRICS

Extinction metrics were generated from PHYLACINE version 1.2.1 (Faurby et al., 2018), a repository of range maps, trait data and phylogeny for all extant and extinct mammal species recorded since 130,000 ybp. The repository includes two classes of maps, the current ranges, and the present natural ranges. The present natural ranges represent the distributions of extinct and extant mammal species if they had never experienced anthropogenic pressures whereas the current ranges include human induced changes to a species’ distribution. The maps are projected to Behrmann cylindrical equal area rasters with a cell size of 96.5 km x 96.5 km at 30° North and 30° South.

A schematic summarizing how the extinction metrics were generated is available in Figure 1. For each extant host species used by Olival et al. (2017) and Shaw et al (2020), the corresponding PHYLACINE present-natural range map was compared to the other present-natural range maps for extinct and other extant mammals. Given the relatively low resolution of the repository maps, sympatry was defined as any distribution overlap. This resulted in two metrics for each Olival et al. (2017) host, a count of sympatric extinct mammals and sympatric extant mammals. The list of extant mammal species was then subset again, to only those with local extinctions in the range of the Olival et al. (2017) host, caused by human activities. This was achieved by subtracting the present-natural range map (no anthropogenic influences) of each extant sympatric mammal to its current range map (with anthropogenic influences).

**Figure 1.**
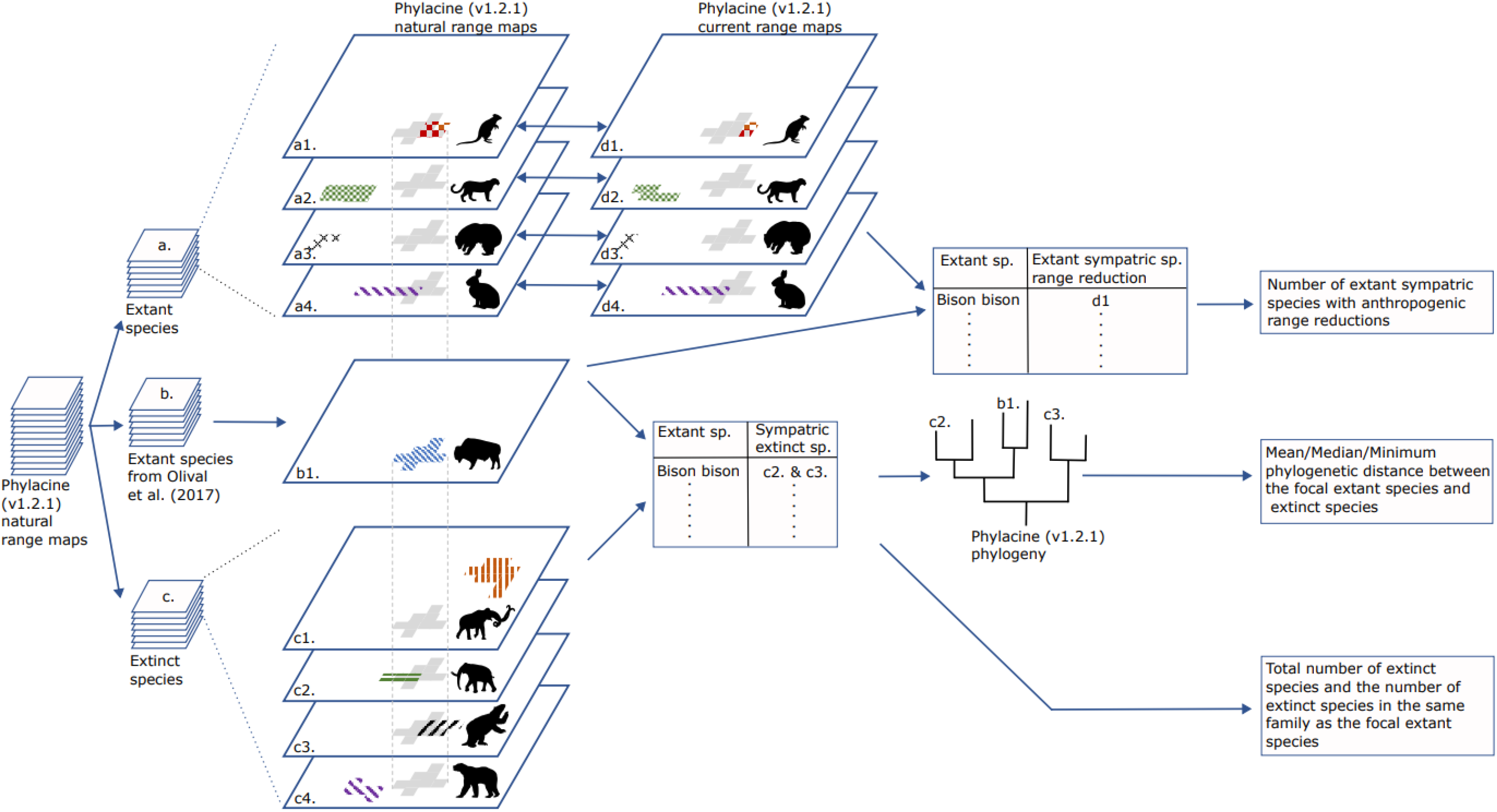
Schematic showing the generation of the extinction variables. All Phylacine range maps (left-hand side) represent extinct and extant species distributions under the assumption of no anthropogenic influence. Range maps were subdivided into the extant hosts (a), of which a subset was included in the Olival et al. (2017) study (b), as well as extinct species (c). Each of the host species from Olival et al. (2017) (b1) were compared to all other extant species (a1 – 4), to identify the number of extant sympatric species assuming no anthropogenic influence. For each sympatric species, the range map without anthropogenic pressure was compared to the corresponding map with anthropogenic pressure, to identify the number of local extinctions of extant species (represented by d1). The focal range map (b1) was also compared to all extinct ranges (c1 – 4), to identify all those that overlapped (c2 & c3). The Phylacine phylogeny was then used to determine the minimum, mean and median phylogenetic distances between the focal extant species (b1) and sympatric extinct species (c2 & c3). From all those that overlapped, the number of extinct species in the same family as the focal extant species were also determined. Animal silhouettes from PhyloPic.

Next, we calculated the pairwise phylogenetic distances between extinct and extant hosts, to test the hypothesis that more related species would share a greater number of parasites. The complete phylogeny available from PHYLACINE was built using a hierarchical Bayesian approach with full details found in Faurby & Svenning, (2015), and includes the posterior distribution of 1000 trees. Each pairwise distance between mammal species was averaged across 1000 trees, as recommended by the Faurby & Svenning (2015) (and *personal communication*), generating a mammal phylogenetic distance matrix. Using this matrix, we calculated the minimum, mean and median distance between each extant Olival et al. (2017) host and the subset of sympatric extinct mammal species.

As well as phylogenetic distance metrics, we included the number of sympatric extinct species from the same family as the extant focal host. This was done in case the number of closely related species is more important than the mean phylogenetic distance. For example, the same mean phylogenetic distance could be achieved from averaging the distance to one close and one distant relative or averaging the distance to two intermediate relatives. However, counting the number of close relatives (i.e., within the same family) would distinguish these two hypothetical scenarios.

### ADDITIONAL METRICS

The original variables from Olival et al. (2017) were also used here, including research effort, phylogenetically corrected body mass, range, and mammal sympatry. The phylogenetic signal was removed from body mass measurements using a phylogenetic eigenvector regression (PVR) (Diniz-Filho et al., 1998, 2012). The authors calculated research effort from the number of disease-related citations per host species, and mammal sympatry was calculated from IUCN range maps, some of which are impacted by anthropogenic disturbances. Different thresholds for range overlap were used by Olival et al. (2017) to determine mammal sympatry (>0%, ≥20%, ≥40%, ≥50%, ≥80%, or 100%) each of which was included here.

We also added two metrics associated with a slow-fast life history, taken from Albery et al. (2022), but originally compiled by Plourde et al. (2017). These were the first two principal components (PC) from an analysis of six life-history traits associated with increased reproduction over survival. These principal components accounted for 85% of the variation in gestation length, litter size, neonate body mass, interbirth interval, age of weaning and age of sexual maturity. Increasing PC values represent a faster life history.

### GENERALIZED ADDITIVE MODELS (GAMS)

We fitted GAMs by adapting previously published models from (Shaw et al., 2020). The models were implemented using the mgcv package version 1.8.39 (Wood, 2011) in R statistical software v.4.1.3 and used smooth spline predictors and automated double penalty smoothing to remove redundant terms from the models. The default smoothing terms included in the full GAM were as follows:

#### a. Full GAM

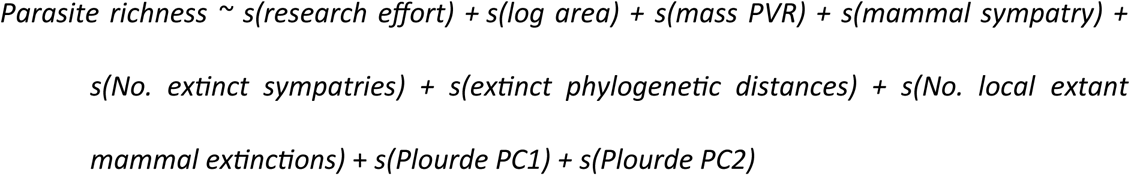

In Equation a., ‘*extinct phylogenetic distances’* represent the minimum, mean or median phylogenetic distance between an extant host and the globally extinct species that were sympatric. Similarly, ‘*No. extinct sympatries*’ represents either the total number of extinct sympatric species, or the number of extinct species within the same family as the extant host. Finally, ‘*mammal sympatry*’ captures each of the thresholds for mammal sympatry between extant species, used by Olival et al. (2017) and described above. In these cases, competing variables for the same mechanistic effect were included in alternative GAMs and the combination of variables resulting in the lowest Akaike Information Criterion (AIC) were selected.

As well as the full GAM model (Equation a.), two reduced models were also fit to the same parasite richness data (Equations b. and c., below). The first had neither the extinction nor fast-life history metrics and allowed us to compare the full models to GAMs fitted using only the variables included by Olival et al. (2017) and Shaw et al. (2020) (Equation b.). Model fit was assessed by comparing the deviance explained and the AIC between Equations a. and b. The second reduced model (Equation c.) only excluded the extinction variables and was used to assess the geographic patterns in pathogen richness due to lost mammal species (described further in section 3.5).

#### b. Reduced GAM 1

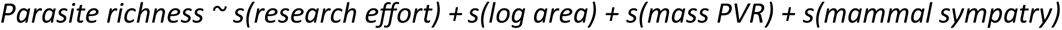

#### c. Reduced GAM 2

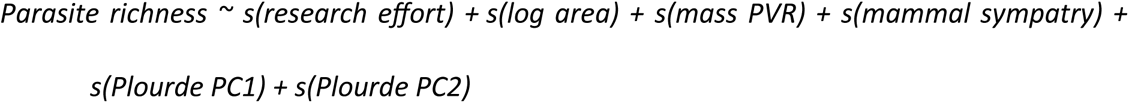

In Equations a-c., the Order of the extant mammal was also fitted as a random effect (not shown). While the above formulas show the default smoothing functions, the formulas were modified when fitting models to bacterial richness data derived from the CLOVER repository (Table S1), due to a lack of interdependence between the smoothing terms as measured by concurvity scores greater than 0.8. This was assessed using the concurvity() function from the mgcv package. In cases of concurvity scores over 0.8, both non-interdependent variables were included in a single smoothing term and alternative GAMs run with each one. Model fit was also assessed using the four plots generated using the gam.check() function available in mgcv: the residual Q-Q plot, histogram of residuals, residuals vs linear predictor plot and observed vs fitted values plot.

### SENSITIVITY ANALYSIS

A sensitivity analysis was performed to assess the impact of adding randomized extinction variables to the reduced model (equation b.) on the AIC score. The aim was to determine if the addition of these variables could produce a similar magnitude decrease in the AIC scores (ΔAIC) as observed between the non-permuted full model and the reduced model (AIC for equation a. minus equation b.). The permutation of variables (sampling without replacement), comparison with the reduced model, and calculation of ΔAIC was repeated 500 times.

### SPATIAL MAPPING OF PARASITE RICHNESS

We used the fitted GAMs to predict the richness of viruses and bacteria for each host species in the (Olival et al., 2017) dataset (n=435 and n=188, respectively), assuming maximum research effort. This involved setting the research effort for each species to the same level as the host species with the highest number of disease-related citations, and then making predictions based on the fitted GAMs. These patterns were visualized geographically using the IUCN (2015.2) range maps, resulting in a raster layer with a resolution of 1/6 of a degree (18km x 18km at the equator). The maps were generated using GAMs fitted with and without extinction variables (described above), such that subtracting one from the other showed regions of the world where pathogen richness is expected to differ due to historical mammal losses. For each of viral and bacterial richness, a map was generated across all mammalian orders as well as individual maps for Carnivora, Cetartiodactyla, Chiroptera, Primates and Rodentia.

## RESULTS

### GAM FITS

Results from GAMs fitted to per-host virus and bacteria richness data from (Shaw et al., 2020) are presented hereafter, with results from the CLOVER per-host parasite richness data presented in the Supplementary Material (Table S2 & Figure S2, S5, S6).

The addition of extinction and fast-life history variables increased the deviance explained and reduced AIC scores compared to models without these metrics. Full GAMs explained 58.8% and 72.9% of the deviance in viral and bacterial richness per host (Table 1), an improvement of 5.4% and 4.9% over models fitted without the metrics (Equation b., above), respectively. AIC scores reduced by 41.91 and 16.34 for GAMs fitted to virus and bacteria richness data (from 1871.01 to 1829.10 and 653.53 to 637.19), respectively.

**Table 1.**
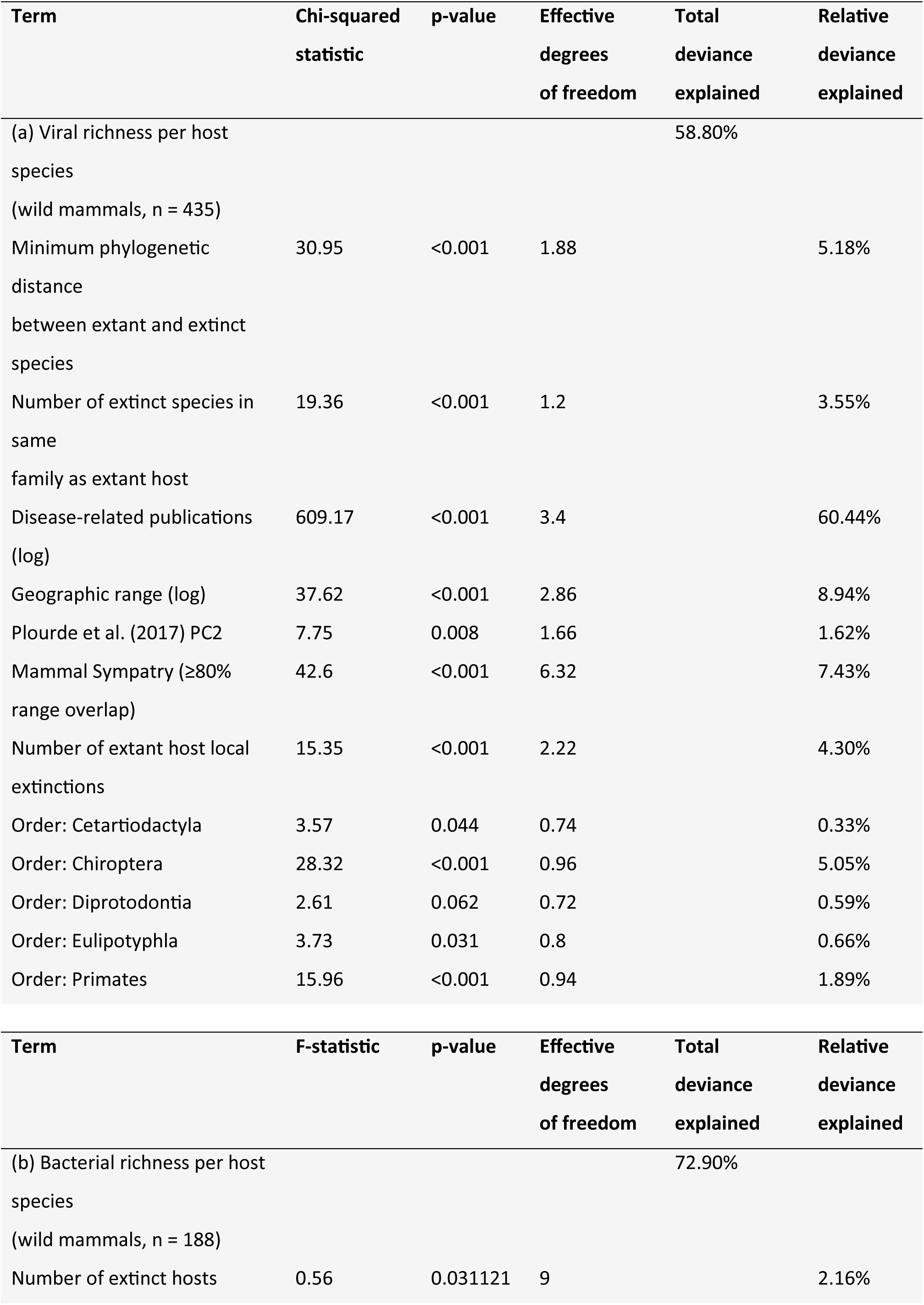
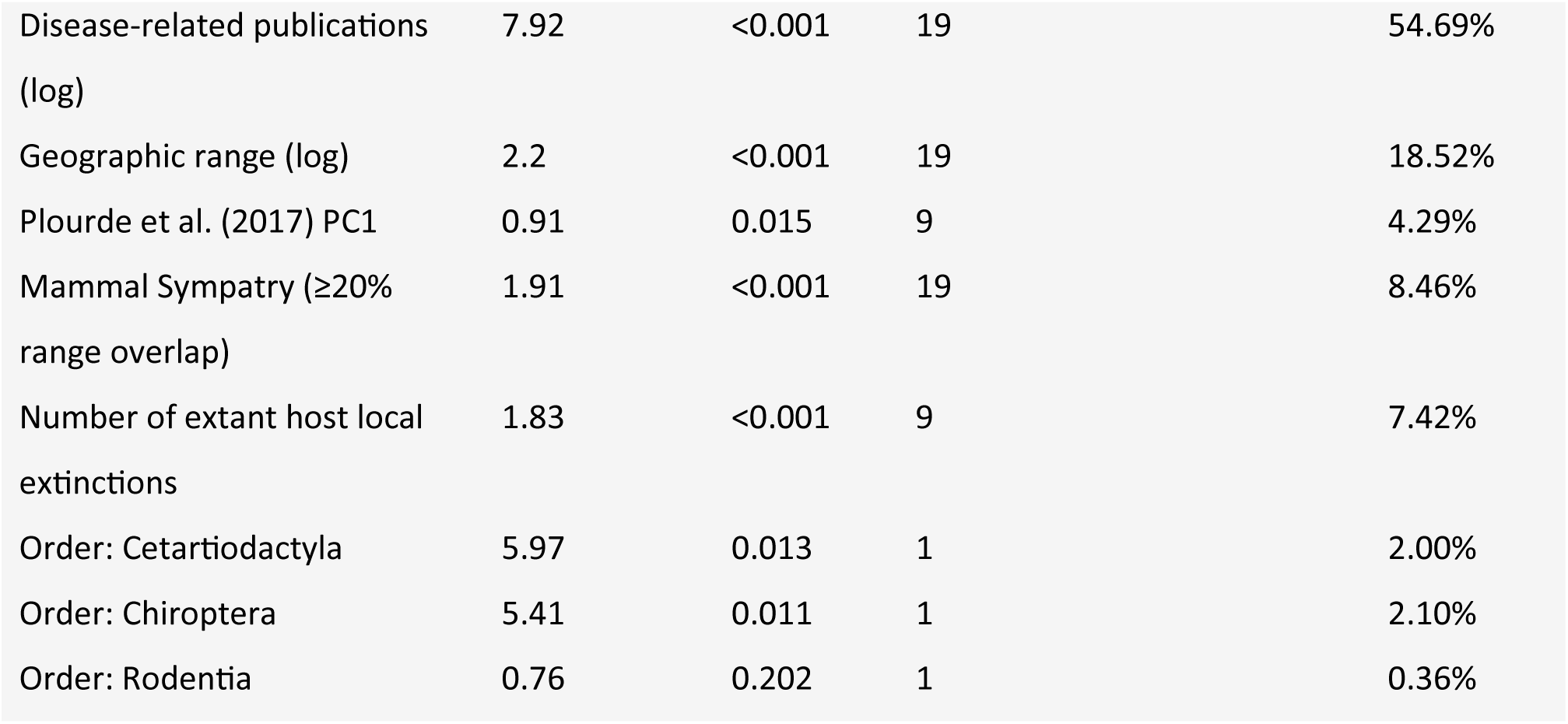
Summary of GAMs fit to per-host viral and bacterial richness data from (Shaw et al., 2020). Only input variables significantly associated with per-host richness are included.

Across all variables, research effort explained the highest relative deviance of per-host viral (60.44%) and bacterial (54.69%) richness, with a significant positive association for each pathogen type (Figure 2c and 2j, respectively; Table 1: p-value <0.001). For viruses, the extinction metrics accounted for a greater relative deviance than any other biologically meaningful variables. Metrics associated with local and global mammal extinctions had a cumulative relative deviance of 13.04% and 9.60% of viral and bacterial richness, respectively. These translated to 32.96% and 21.14% of the relative deviance explained by biologically meaningful variables (all variables bar research effort), respectively. For bacteria, the relative deviance for extinction metrics fell behind that of host geographic range.

**Figure 2.**
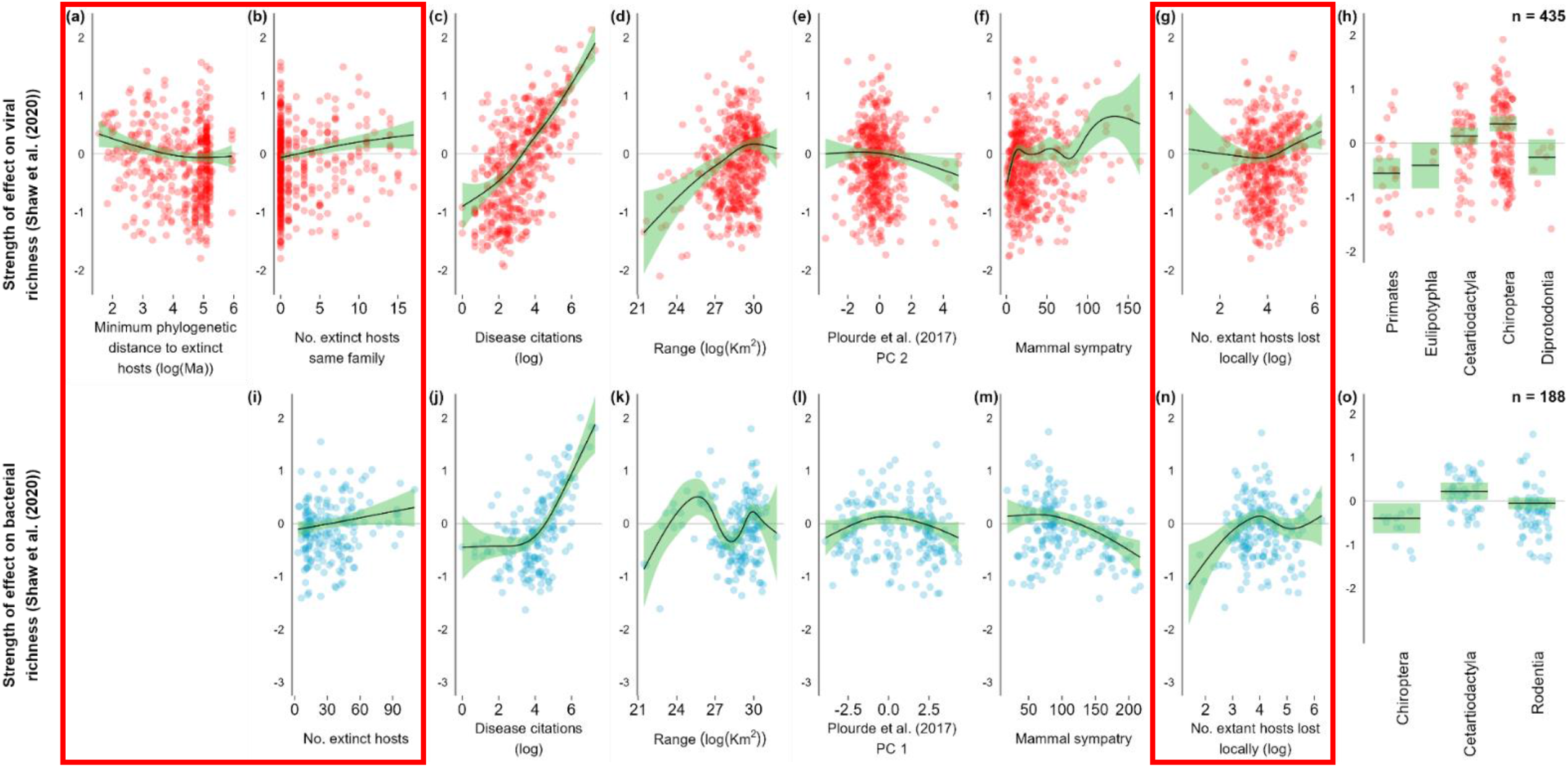
Partial dependence plots for GAMs fitted to viral (top), and bacterial (bottom) richness in extant mammalian species. Data on host-viral and -bacterial associations were taken from the Shaw et al. (2020) dataset. The GAMs were initially formulated by Olival et al. (2017), were subsequently used by Shaw et al. (2020), and included many of the same variables, including disease citations, host range and mammal sympatry. To these were added novel extinction variables associated with Phylacine (version 1.2.1), summarized in Figure 1. Partial dependence plots for these novel extinction variables are outlined in red. Additionally, the first two mass-corrected principal components from Plourde et al. (2017) were also incorporated, which represent 85% of variation across 6 fast-slow life history traits: gestation length, litter size, neonate body mass, interbirth interval, weaning age, and sexual maturity age.

The sensitivity analysis compared the ΔAIC between the reduced GAM in equation b. and the same GAM either with the extinction variables or permuted extinction variables (number of repeats: 500). For viral and bacterial richness, 3% and 9% of GAMs with permuted extinction variables had an equivalent or lower AIC than the GAM with non-permuted extinction variables (Figure S3a and S3b), respectively.

Partial dependence plots show that viral richness tends to increase with the number of extinct hosts in the same family as the focal extant host (Figure 2b; Table 1: p-value <0.001) but decreases as minimum phylogenetic distance increases (Figure 2a; Table 1: p-value <0.001). The local extinction of extant sympatric mammals is also associated with greater viral richness in hosts (Figure 2g; Table 1: p-value <0.001). Like viruses, the extinction variables were significantly associated with bacterial richness. Local and global extinctions of sympatric mammal species were both positively associated with bacterial richness (Figure 2n and Figure 2i, respectively; Table 1: p-value <0.001 and 0.031, respectively). Although model fit was less favorable when fitting GAMs to CLOVER data, the partial dependence plots revealed broadly similar patterns between the extinction variables and pathogen richness (Figure S2).

Mammal range was significantly associated with an increasing per-host viral richness (Figure 2d; Table 1: p-value <0.001, respectively), but displayed a relationship with bacterial richness containing multiple inflexion points (Figure 2k; Table 1: p-value <0.001). Mammal sympatry was also significantly associated with increasing per-host viral richness (Figure 2f; Table 1: p-value <0.001) but decreasing bacterial richness (Figure 2m; Table 1: p-value <0.001). PC2 from Plourde et al. (2017) had a slight but significant negative association with viral richness (Figure 2e; Table 1: p-value = 0.008) whereas PC1 had an inverted U-shaped relationship with bacterial richness (Figure 2l; Table 1: p-value = 0.015). Mammalian orders associated with increased numbers of virus species per host included Chiroptera and Cetartiodactyla (Figure 2h; Table 1: p-value <0.001 and 0.044, respectively), with the latter taxon also associated with greater bacterial richness (Figure 2o; Table 1: p-value = 0.013).

### GEOGRAPHIC PATTERNS OF PATHOGEN RICHNESS DUE TO LATE PLEISTOCENE EXTINCTIONS

Subtracting pathogen richness maps generated with extinction metrics from maps generated without these variables (Figure S4) shows that the impact of the extinctions was geographically heterogeneous (Figure 3). Broadly, the inclusion of these metrics results in higher levels of viral and bacterial richness in South and North America (Figure 3a and 3b), with virus richness also elevated in Eurasia and Oceania.

**Figure 3.**
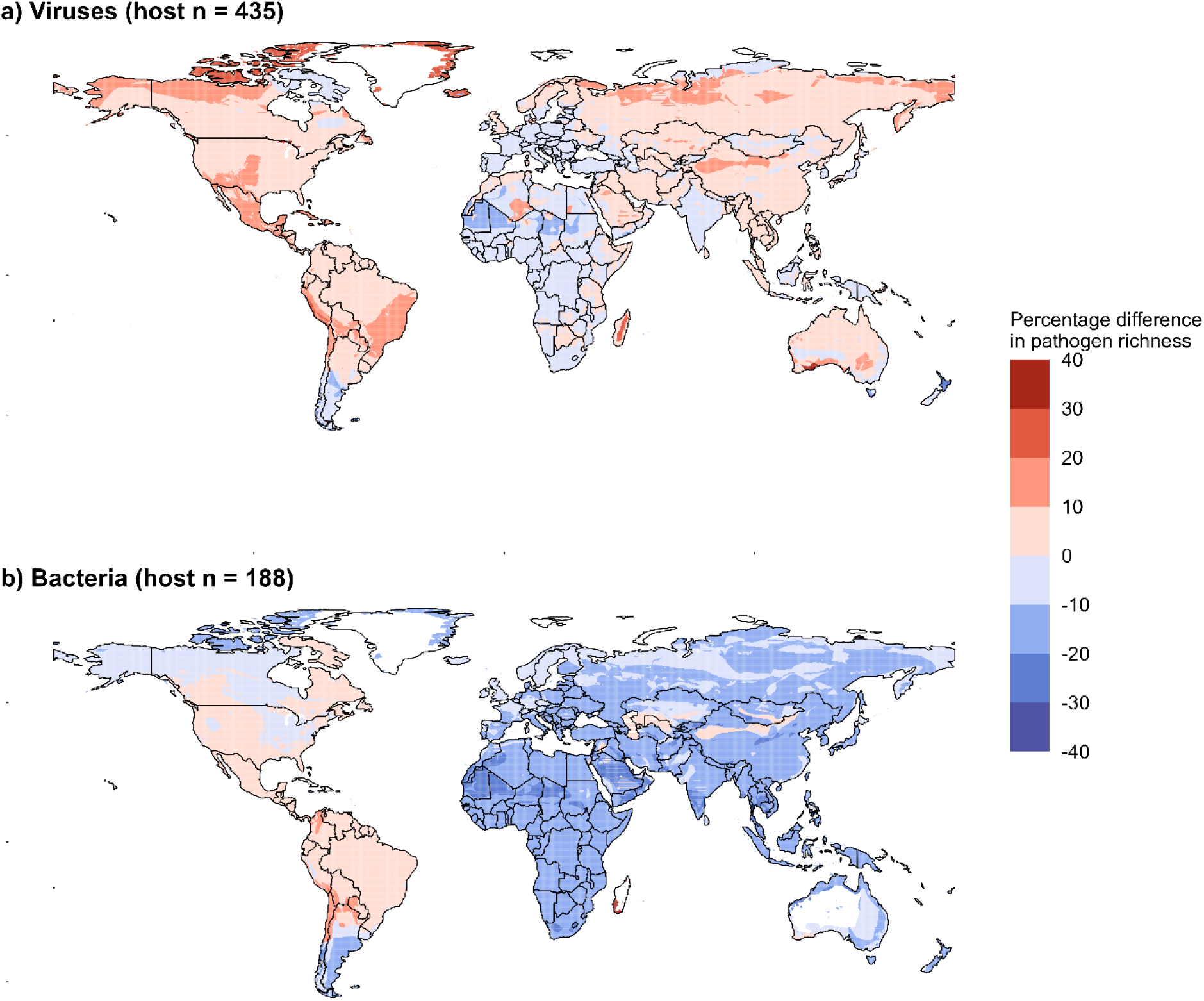
Maps showing the change in geographic patterns of virus (a), and bacteria (b) richness due to historical mammal extinctions. Viral and bacterial richness was calculated for host species using two generalized additive models (GAMs) (both assuming maximum research effort). One class of models included variables related to historical mammal losses and the other class did not. The results from both models were visualized using IUCN range maps (2015.2), and the map generated without extinction variables was subtracted from the map that did, resulting in maps a and b.

Areas of increased viral richness included all continents bar Africa, which had −2.6% fewer viruses. Higher predicted viral counts were observed in North America (8.6%), South America (6.8%), Asia (3.4%), Europe (5.1%) and Oceania (3.3%) (Figure 3a). Lower viral richness in Africa was common to each of the mammalian families Carnivora, Chiroptera, Primates, and Rodentia (Figure S5a, c, d, and e, respectively), but not Cetartiodactyla (Figure S5b). Host species from Carnivora and Rodentia were also predicted to have increased viral richness in Eurasia, whereas Cetartiodactyla occupying a central region of Asia were predicted to have a higher viral richness.

The extinctions also lead to a higher predicted bacterial richness in South America (3.1%), with all other continents showing decreased richness (North America: −1.8%; Europe: −10.2%; Oceania: −11.6%; Asia: −13.6%; Africa: −15.4%) (Figure 3). When separated by mammalian order, areas of increased bacterial richness (due to the extinction metrics) included much of South America for Carnivora, North and Central America for Cetartiodactyla (Figures S6a and S6b, respectively), and regions of southeast Asia, east Africa, and South America for Primates (Figure S6d).

## DISCUSSION

Phylogenetic proximity and mammal sympatry have previously been identified as variables positively associated with viral sharing (Albery et al., 2020; Cooper et al., 2012; Davies & Pedersen, 2008; Parrish et al., 2008) and per-host pathogen richness (Olival et al., 2017; Shaw et al., 2020), respectively. Shorter phylogenetic distances increasing parasite sharing between hosts is common across the tree of life (Longdon et al., 2014), from rabies virus in bats (Streicker et al., 2010), nematodes and RNA viruses in fruit flies (Longdon et al., 2011; Perlman & Jaenike, 2003), to fungal pathogens infecting plant species (Gilbert & Webb, 2007). While phylogeny determines the maximum number of hosts a pathogen can infect, the actual host range is a subset of this theoretical maximum and is affected by whether the pathogen is presented with an opportunity to contact a novel host (Perlman & Jaenike, 2003). Thus, mammal sympatry may limit the number of hosts utilized by a parasite.

Results presented here indicate that phylogenetic distance and mammal sympatry should also be considered between extant and extinct hosts in future studies modelling per-host pathogen richness. We show that extant mammal species overlapping in their range with many phylogenetically proximate hosts tend to have a greater richness of viruses and bacteria. Our results lead us to ask, why historical mammal losses are associated with an increased richness of pathogens in extant hosts? We list four non-mutually exclusive mechanisms below for why this might be, illustrated in Figure 4.

**Figure 4.**
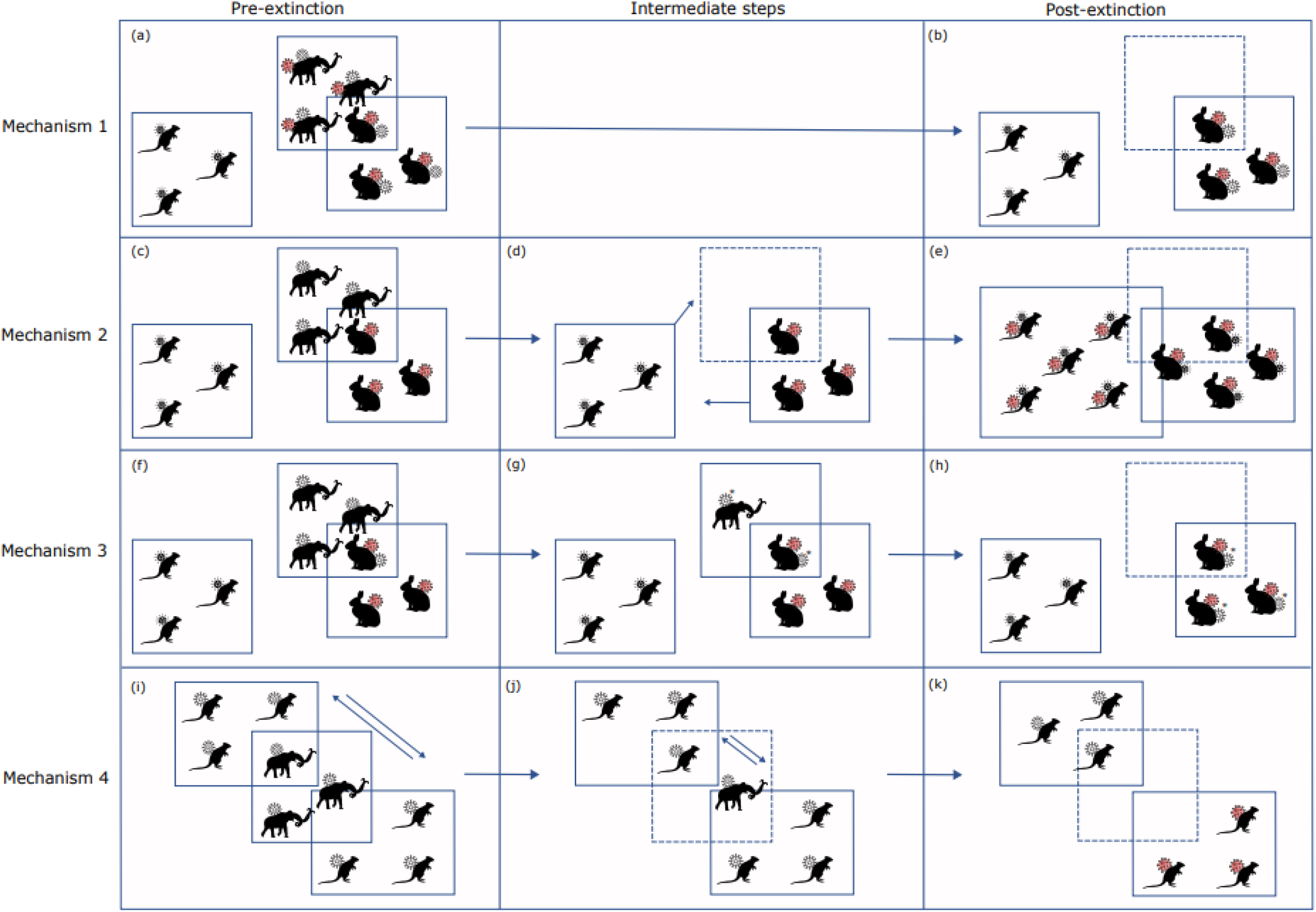
Illustrates five mechanisms for how megafauna extinctions could be correlated with increased parasite richness in surviving hosts. The left-hand column shows the distributions (internal boxes) of three species prior to an extinction event, which removes the mammoth species. The pathogens infecting these species are represented as virions contacting each host outline. The two rodent schematics represent extant hosts that persist after extinction with a greater pathogen richness (right-hand column). In Mechanism 1 (first row) pathogen and host richness are correlated prior to the extinction of the mammoth species (a). Following the mammoth extinction, its pathogens persist in the sympatric extant species (b). Mechanism 2 (second row) illustrates how the extinction of the megafauna species has a cascading effect in the ecosystem (c to d), such that new contacts are made between the surviving species, allowing for novel pathogen transmission (e). Mechanism 3 (third row) shows how the mammoth pathogen is capable of infecting sympatric mammals at a low level (f). However, during the extinction event (g) there is selection on the pathogen (*) to adapt to the sympatric extant host, enabling it to circulate in that host after the global extinction of the mammoth (h). Mechanism 4 shows how the removal of the mammoth (i to j) reduces the spread of pathogens across the landscape, leading to their divergence (k; illustrated by the change in virion color).

### Mechanism 1 (*Figure 4*)

*A portion of the parasites infecting mammal species destined for extinction (since the Late Pleistocene) were able to persist in surviving hosts*. This would suggest that the extinctions left behind hosts with a pathogen richness now disproportionate to the number of remaining sympatric species. While Dunn et al. (2010) show a positive correlation between pathogen and extant host richness, pathogen richness in any given area may better reflect the number of extinct plus extant host species.

Few studies to date have recorded the co-extinction of parasites following host extinctions, which may reflect the difficulty in detecting these events, or that parasites have a greater capacity to utilize other hosts in their environment than has been previously appreciated (Dunn et al., 2009). Of the parasites infecting now-extinct mammals, the true specialists, and generalists unable to maintain a NRR > 1 with the demise of the primary host would likely have faced co-extinction (Farrell et al., 2015), whereas generalists capable of an NRR > 1 may have persisted. However, a parasite’s true host specificity may differ from the actual observed specificity today, making it difficult to predict whether they would be threatened with co-extinction at the demise of their primary host. Parasites that appear as specialists in the present day may once have been generalists, also capable of infecting the lost species prior to their extinction. Farrell et al. (2021) show an increase in the evolutionary distinctiveness of extant host species following the extinctions, limiting the opportunities for their parasites to move into a phylogenetically proximate host, making them appear as specialists.

Interestingly, a study by Walker et al. (2017) showed that larger hosts tend to carry a greater number of generalist parasites. This would be significant for the species lost during the Late Pleistocene, which were among the largest terrestrial mammals (Smith et al., 2018), suggesting that many of their pathogens may have been candidates to survive in extant mammal species. However, this relationship between host size and the specificity of the cargo parasites is not clear, as Krasnov et al. (2006) show that in certain systems larger species tend to carry specialists.

### Mechanism 2 (*Figure 4*)

*Host extinctions caused cascading effects in the ecosystem, leading to novel contact between extant species and transmission of their pathogens.* Here we propose that the loss of mammals during the Late Pleistocene and early Holocene changed the assemblage and interaction patterns amongst the surviving host species (Galetti et al., 2018). Their removal would leave niches available for surviving hosts to move into. Additionally, larger species provide many services, including modifying the physical structure of an ecosystem as well as the dispersal of nutrients and seeds (Malhi et al., 2016). Therefore, the extinctions modified multiple environmental features, potentially changing contact patterns between extant hosts, and facilitating novel pathogen transmission.

### Mechanism 3 (*Figure 4*)

*Parasites carried by species destined for extinction adapt to novel hosts.* Theory suggests that pathogens face a trade-off between infecting a greater number of hosts with reduced virulence per host, or fewer hosts with increased virulence in each one (Dunn et al., 2009). In many cases a parasite must adapt to infect, and thereafter be transmitted by, a novel host (McCarthy et al., 2007; Poullain and Nuismer, 2012; Vanderford et al., 2007). These evolutionary changes may come at a cost to other aspects of the parasite’s reproductive success, including its ability to replicate in its original host (Ebert, 1998). However, reduced infectivity in the original host would not be a real cost if this species was destined for extinction. Cooper and Scott (2001) showed that serial passage of an arbovirus in cells from either of its original hosts resulted in adaptive evolution to that species, highlighting the pathogen’s ability to evolve to the available host. The parasite’s success in adapting to a new host would likely depend on the existing genetic variation within the parasite population and the speed of host extinctions, which determine the available time for the required adaptations to evolve.

### Mechanism 4 (*Figure 4*)

*Loss of the larger mammal species and their dispersal services would disconnect subpopulations of pathogens, leading to their diversification*. Doughty et al. (2020) estimate a ∼85% reduction in the dispersal of generalist microbes and ectoparasites carried by terrestrial hosts following the extinctions that occurred during the Late Pleistocene and early Holocene. While there is limited empirical evidence of microbe diversification as a direct result of megafauna extinction, support for this mechanism comes from the theory of island biogeography (Heaney, 2000). Bergeijk et al. (2022) recently showed the high degree of phylogenetic and metabolic dissimilarity between Actinobacteria from the gut of a 28,000-year-old mammoth and modern members of this bacterial phylum. The authors highlight that these differences may be due to a low number of reference genomes available in certain Actinobacteria taxa. As these genomes become available, it may be possible to determine whether the genetic difference between the mammoth and modern Actinobacteria species is disproportionate to their age difference, which would support this mechanism.

Our results suggest that host losses can have long-lasting effects on parasite richness, potentially persisting for tens of thousands of years. While we have identified a positive relationship between pathogen richness and hosts lost since the Late Pleistocene, these patterns may be obscured in the coming years, as human actions continue to remove mammal species and shift their distributions (Carlson et al., 2021). Anthropogenic pressures are driving a sixth mass extinction (Pievani, 2014), and the influence of these new host losses may be detectable as changes in pathogen richness for millennia to come.

Maps generated in this study show that viral and bacterial richness is marginally higher in areas that lost greater numbers of Late Pleistocene mammals. The Americas lost a particularly high number of megafauna species, including multiple species of mammoth, mastodons, and ground sloths (Stuart, 2015), and show that these areas have a higher predicted richness of viruses and bacteria. Increased viral richness (due to the extinction variables) is also observed in Europe, Oceania, and Asia, which also lost many mammal species during the Late Pleistocene and early Holocene. Hence, the inclusion of extinction variables in our models suggest that these areas should receive increased surveillance for viruses and bacteria compared to surveillance efforts based on models run without these metrics.

The PCs that captured variation across the six fast-slow life history variables were not correlated with a clear increase in pathogen richness for either virus or bacterial richness, in contrast to a recent study by Albery et al. (2022). The authors here showed that a faster life-history (increasing PC values) is correlated with a greater viral richness, however here we show that higher PC2 values and extreme PC1 values are associated with lower virus and bacterial richness, respectively.

Inconsistencies between the results of this study and previous literature may be due to the low proportion of total host-pathogen associations currently identified. We refer readers to Olival et al. (2017) for maps showing the distribution of mammal species with uncharacterized viruses. This is not necessarily a limitation if the relative pathogen richness between hosts remains stable with increasing sampling. However, Gibb et al. (2022) show that the per-host virus diversity across mammals is highly sensitive to short term changes in research effort, and the authors err caution when interpreting results from studies using these datasets. This result led us to compare the relationship between input variables and pathogen richness when using all known host-pathogen associations (CLOVER repository) or a subset of these compiled by systematic literature review only (Shaw et al. (2020)). We found that both datasets showed similar relationships between pathogen richness and the novel extinction variables, supporting their inclusion in future studies. However other variables were not identified across both datasets. GAMs run on the CLOVER dataset did not identify associations between bacterial richness and host range, or viral richness and PC2 from Plourde et al. (2017), suggesting that these relationships need further evaluation with datasets containing a greater number of host-pathogen associations.

Despite the limited resolution of the Phylacine range maps, a link was found between the extinction of host species and increased pathogen diversity in the surviving hosts. However, improvements to the resolution of these maps through aDNA techniques may elicit a stronger association. Increasing numbers of studies characterize aDNA from sediments (sedaDNA), removing the need for a paleontological specimen to infer the species’ range (Haile et al., 2009).

The lack of pathogen data from extinct species is also a limitation. Here we use mammal sympatry and phylogenetic distance to extinct mammals to explain the pathogen richness in extant species, but we do not use pathogen data to connect these two cohorts of hosts (e.g., in a bipartite network). However, recent advances in techniques for classifying highly degraded aDNA (Jensen et al., 2019) may allow us to measure the pathogens of extinct late Pleistocene mammals directly and collate databases of ancient host-pathogen associations. This will be important in distinguishing between the different mechanisms (outlined above) driving our finding, that extant mammals tend to have a greater pathogen richness when their ranges overlap with more phylogenetically similar hosts. Understanding the long-term implications of host extinctions is needed currently, as anthropogenic pressures continue to threaten a growing proportion of wild species (Pievani, 2014), with many extinctions projected in coming years.

