## Supplementary Material for "The impact of late Pleistocene mammal extinctions on pathogen richness in extant hosts"

Address: 70d Highgate West Hill, London, N6 6NU

**Keywords:** Host-pathogen associations, zoonoses spillover, emergent infectious disease, Late Pleistocene extinctions

### SUPPLEMENTARY MATERIAL

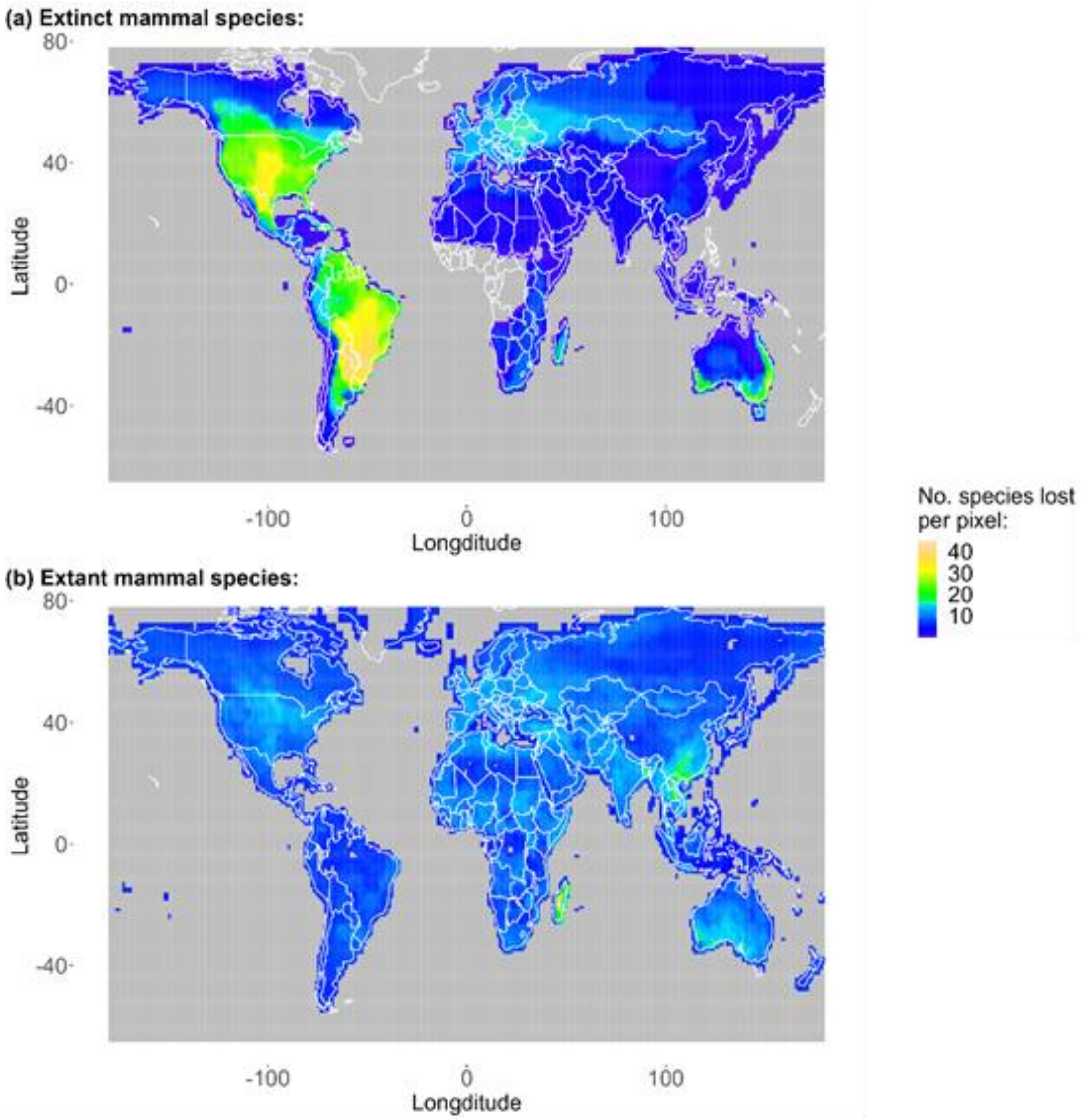


**Figure S1.** Maps generated from Phylacine (version 1.2.1) range maps showing the geographic richness of extinct mammal species (a; present natural ranges) and the number of extant mammal species that are estimated to have been lost per pixel (local extinctions) due to anthropogenic influences (b; present natural – current range maps), recorded since 130,000 ybp. Map (a) highlights hotspots of extinction in North and South America with no recorded mammal extinctions from central western Africa.

| **Pathogen type** | **Repository** | **GAM formula** | **GAM family** |
|---|---|---|---|
| Virus | Shaw et al. (2020) | *viral richness* ~ s(research effort) + s(*log area*) + s(m*ass PVR*) + s(*mammal sympatry*) + s(*extinct sympatry*) + s(*extinct phylogenetic distance*) + s(*Plourde PC1*) + s(*Plourde PC2*) + s(*number of local extant mammal extinctions*) | Poisson |
| Bacteria | Shaw et al. (2020) | *viral richness* ~ s(research effort) + s(*log area*) + s(m*ass PVR*) + s(*mammal sympatry*) + s(*extinct sympatry*) + s(*extinct phylogenetic distance*) + s(*Plourde PC1*) + s(*Plourde PC2*) + s(*number of local extant mammal extinctions*) | Tweedie |
| Virus | CLOVER | *viral richness* ~ s(research effort) + s(*log area*) + s(m*ass PVR*) + s(*mammal sympatry*) + s(*extinct sympatry*) + s(*extinct phylogenetic distance*) + s(*Plourde PC1*) + s(*Plourde PC2*) + s(*number of local extant mammal extinctions*) | Tweedie |
| Bacteria | CLOVER | *viral richness* ~ s(research effort) + s(*log area*) + s(m*ass PVR*) + s(*mammal sympatry*) + **s(*extinct sympatry OR extinct phylogenetic distance**)** + s(*Plourde PC1*) + s(*Plourde PC2*) + s(*number of local extant mammal extinctions*) | Tweedie |

*Just the minimum phylogenetic distance between extinct and extant species.

**Table S1**. Showing the formulas of the GAMs best fit to pathogen richness data from the CLOVER and Shaw et al. (2017) datasets. Many of the smoothing terms include general names that cover several different variables. As in Olival et al. (2017), mammal sympatry represents a range overlap of either >0%, ≥20%, ≥40%, ≥ 50%, ≥80%, or 100%, whereas extinct sympatry represents either the number of extinct hosts lost from the range of an extant mammal species or the number of extinct hosts from the same family as the extant host that were lost from that range. Extinct phylogenetic distance represents either the median, mean, or minimum phylogenetic distance between the extant host and the subset of extinct species that were once sympatric. For bacterial richness data from the CLOVER repository, the extinct sympatry and extinct phylogenetic distance metrics were included in a single smooth term (also emboldened), due to concurvity scores ≥0.8 these variables were included separately.

| Term | F statistic | p-value | Effective degrees of freedom | Total deviance explained | Relative deviance explained |
|---|---|---|---|---|---|
| (a) Viral richness per host species (wild mammals, n = 511; CLOVER dataset) | | | | 54.10% | |
| Median phylogenetic distance between extant and extinct species | 1.64 | <0.001 | 1.95 | | 3.59% |
| Number of extinct species in same family as extant host | 0.37 | 0.103 | 0.67 | | 0.63% |
| Disease-related publications (log) | 22.17 | <0.001 | 3.68 | | 80.40% |
| Geographic range (log) | 0.3 | 0.0725 | 0.81 | | 0.62% |
| Mammal Sympatry (≥40% range overlap) | 1.92 | 0.002 | 2.78 | | 3.51% |
| Non-anthropogenic mammal sympatry | 2.61 | <0.001 | 1.76 | | 3.93% |
| Order: Eulipotyphla | 10.54 | <0.001 | 0.91 | | 1.85% |
| Order: Primates | 16.01 | <0.001 | 0.94 | | 3.90% |
| Order: Rodentia | 7.1 | 0.004 | 0.86 | | 1.57% |

| Term | F statistic | p-value | Effective degrees of freedom | Total deviance explained | Relative deviance explained |
|---|---|---|---|---|---|
| (b) Bacterial richness per host species (wild mammals, n = 241; CLOVER dataset) | | | | 64.90% | |
| Number of extinct hosts | 1.52 | 0.051 | 2.38 | | 4.55% |
| Disease-related publications (log) | 8.89 | <0.001 | 3.59 | | 55.62% |
| Plourde et al. (2017) PC1 | 4.66 | <0.001 | 4.18 | | 16.40% |
| Plourde et al. (2017) PC2 | 0.18 | 0.116 | 0.7 | | 0.67% |
| Mammal Sympatry (≥20% range overlap) | 5.24 | <0.001 | 1.99 | | 11.43% |
| Non-anthropogenic mammal sympatry | 1.2 | 0.003 | 1.47 | | 3.60% |
| Order: Cetartiodactyla | 19.21 | <0.001 | 0.92 | | 3.08% |
| Order: Chiroptera | 9.49 | 0.002 | 0.9 | | 2.86% |

| | | | | |
|---|---|---|---|---|
| Order: Rodentia | 7.54 | 0.016 | 0.83 | 1.11% |
| Order: Primates | 2.35 | 0.117 | 0.58 | 0.68% |

**Table S2**. Summary of GAMs fit to per-host viral and bacterial richness data from the CLOVER repository. Only input variables significantly associated with per-host richness are included.

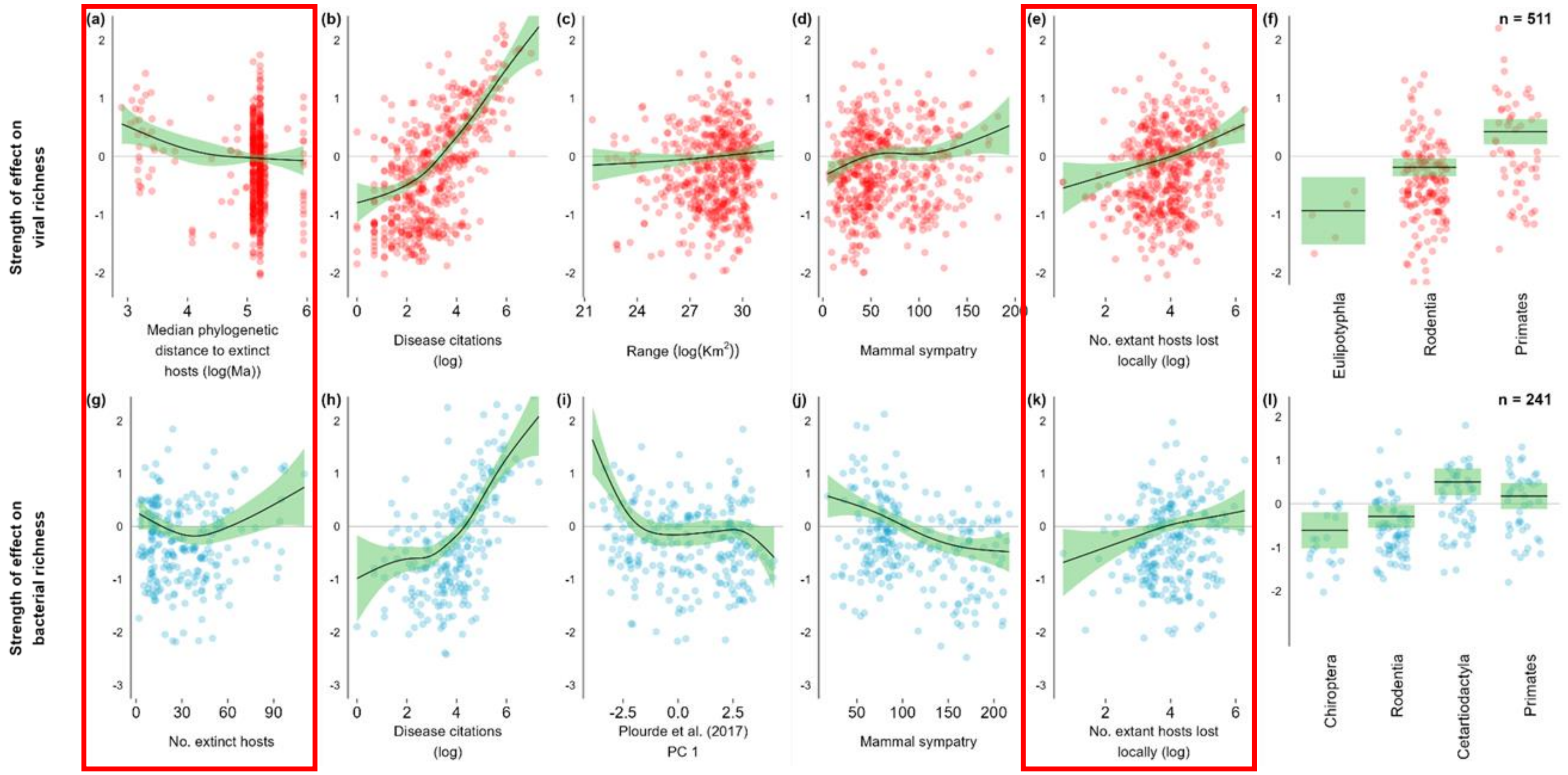


**Figure S2**. Partial dependence plot for GAMs fitted to viral (top) and bacterial (lower) pathogen richness in extant mammalian species, with host-pathogen associations taken from the CLOVER repository. The GAMs were initially formulated by Olival et al. (2017), were subsequently used by Shaw et al. (2020), and included many of the same variables, including disease citations, host range and mammal sympatry. To these were added novel extinction variables associated with Phylacine (version 1.2.1), summarized in Figure 1. Partial dependence plots for these novel extinction variables are outlined in red. Additionally, the first two mass-corrected principal components from Plourde et al. (2017) were also incorporated, which represent 85% of variation across 6 fast-slow life history traits.

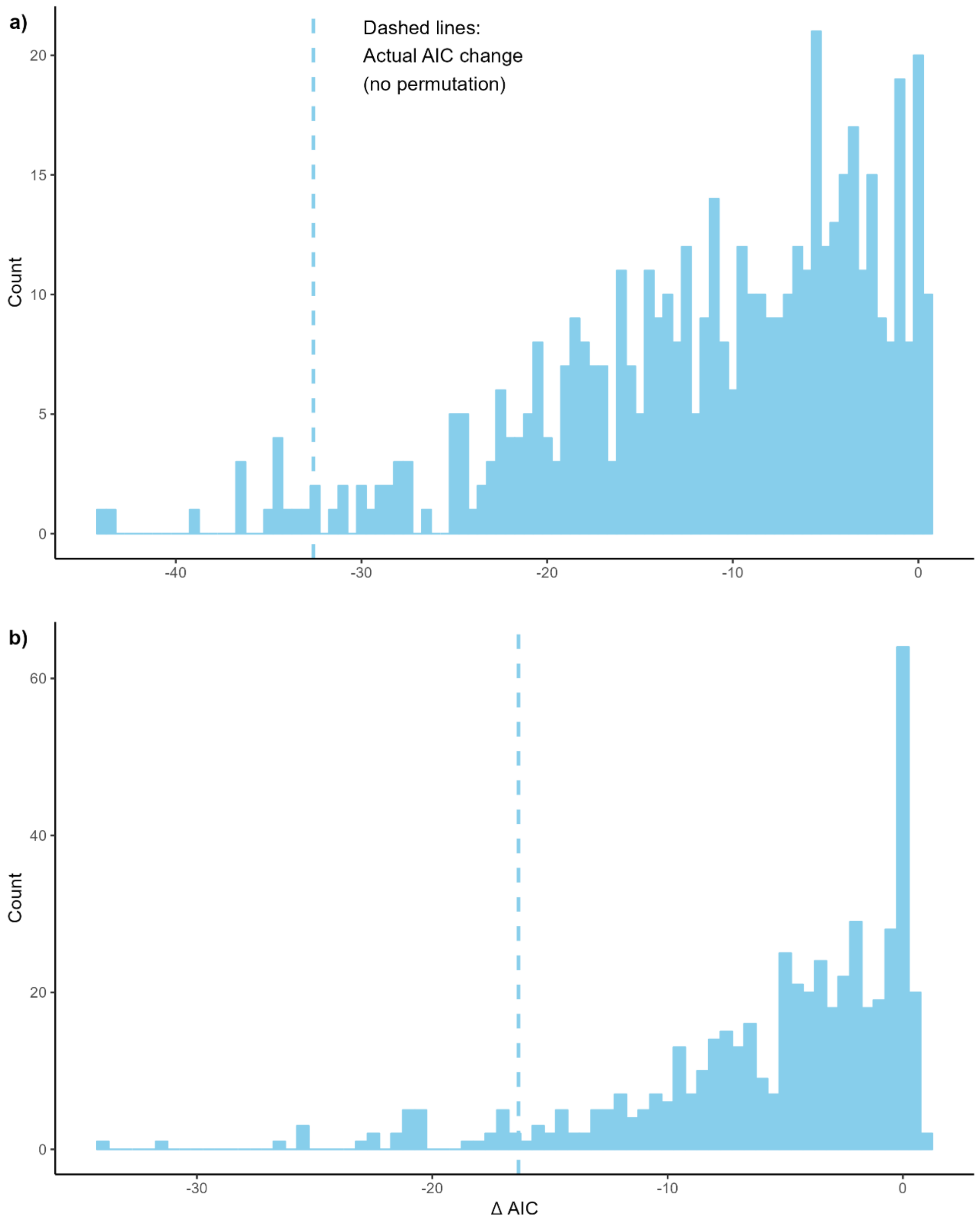


**Figure S3**. Histograms showing the change in AIC scores (ΔAIC) when subtracting the GAM from equation b. (in manuscript) with versus without permuted extinction variables. This calculation was repeated for 500 different permutations of the extinction variables for viruses (upper) and bacteria (lower). This distribution of ΔAIC was compared to the true, observed change in AIC (dashed line), generated by subtracting the score from the full, non-permuted model with the score from the reduced model.

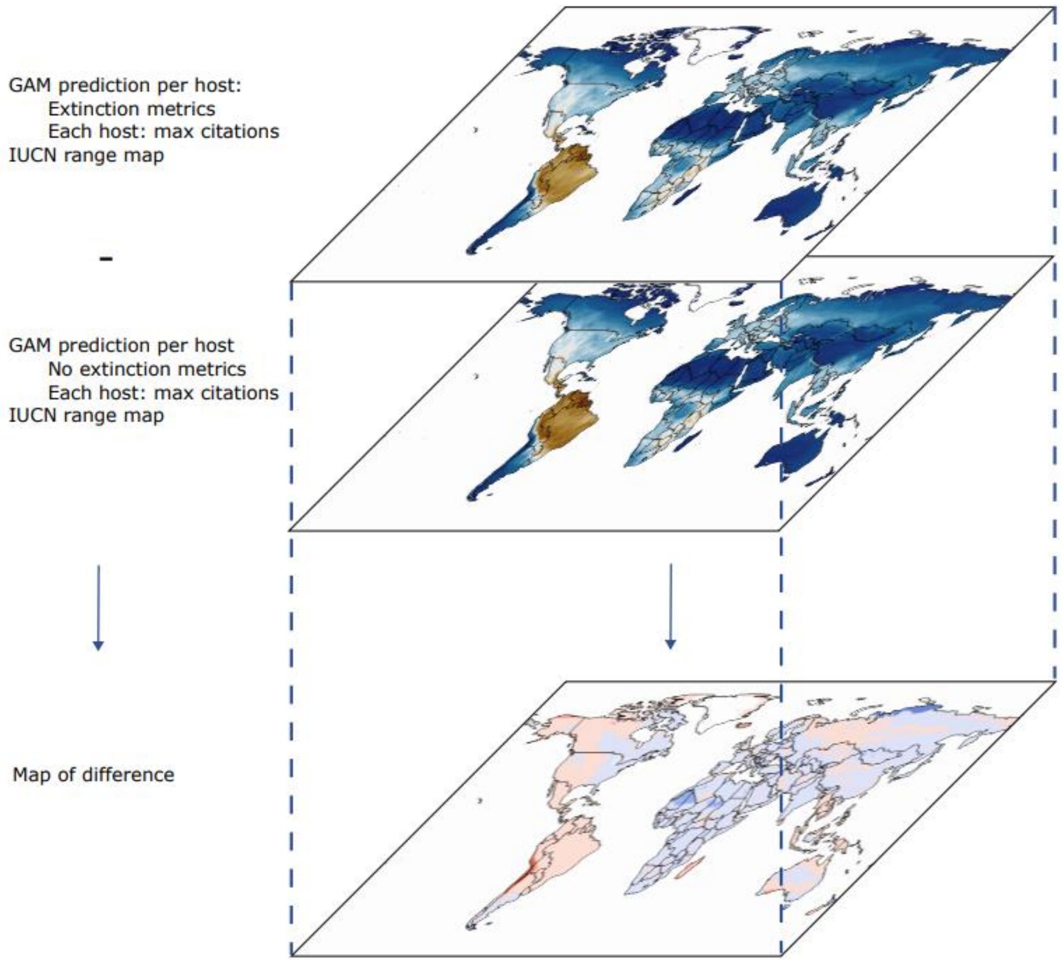


**Figure S4.** Schematic to show the process of map creation. Two maps predicting pathogen richness were created. The first included the extinction metrics (Equation a in manuscript), whereas the second did not include these variables (Equation b.). The two maps were then subtracted from one another, to identify geographic regions where expected pathogen richness changed due to the inclusion of extinction variables.

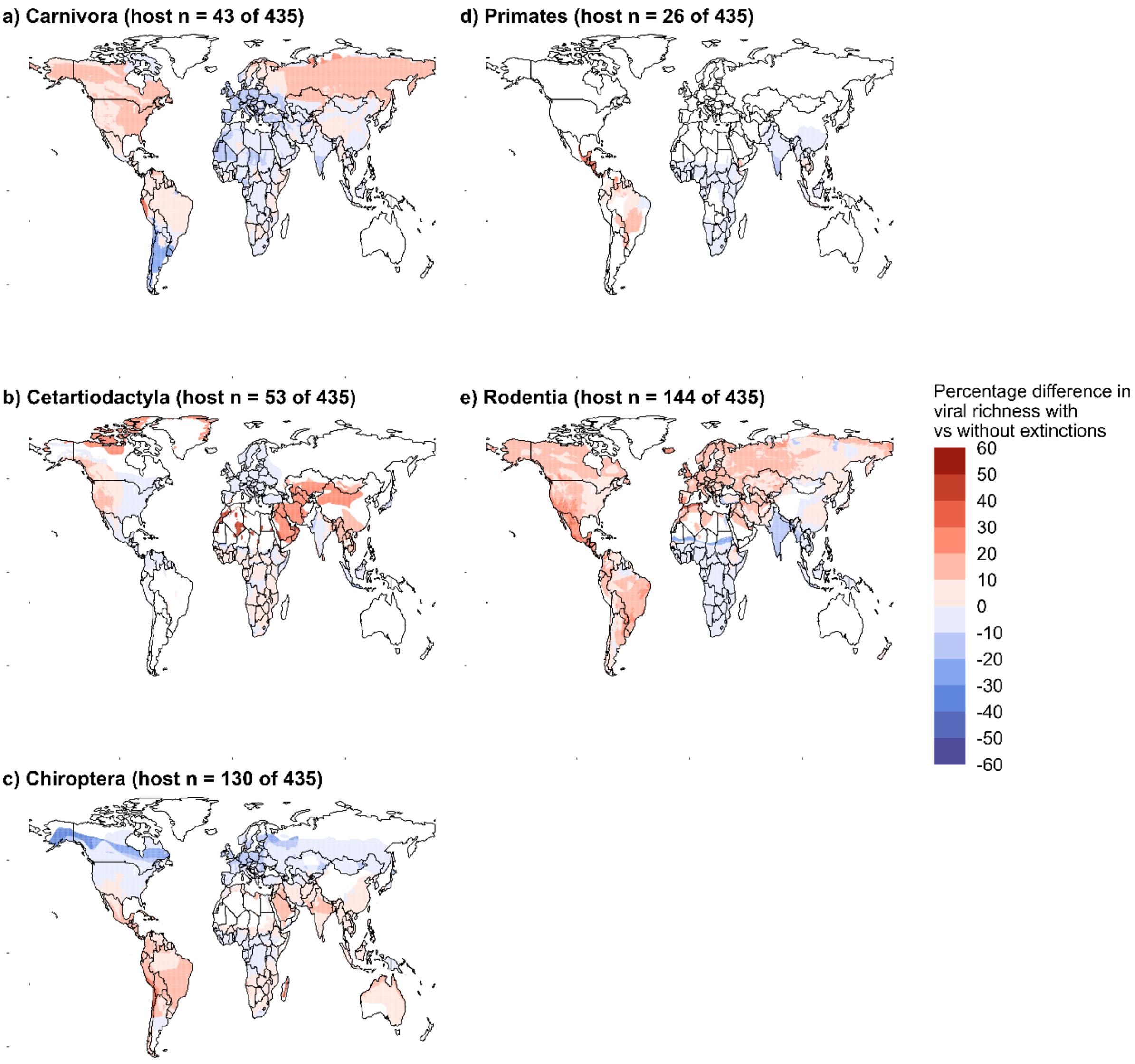


**Figure S5**. Maps showing the change in geographic patterns of **viral richness** caused by historical extinctions when subdividing the extant mammalian hosts by order: Carnivora (a), Cetartiodactyla (b), Chiroptera (c), Primates (d), and Rodentia (e). Viral richness was calculated for each host order using two GAM models (both assuming maximum research effort). One class of models included variables related to historical mammal losses and the other class did not. The results from both models were visualized using IUCN range maps (2015.2), and for each mammalian order, the map generated without extinction variables was subtracted from the map that did, resulting in maps a – c.

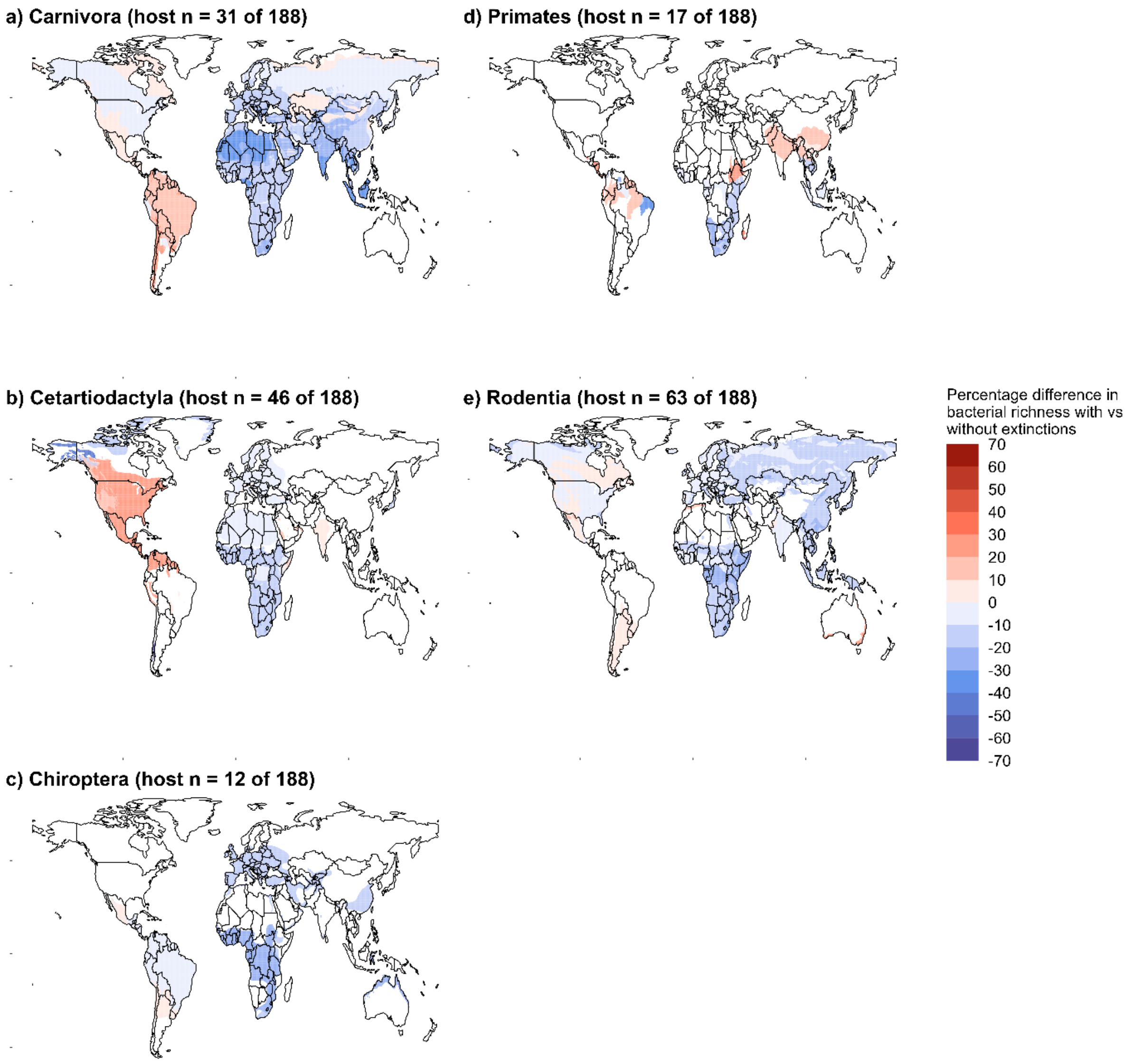


**Figure S6**. Maps showing the change in geographic patterns of **bacterial richness** caused by historical extinctions when subdividing the extant mammalian hosts by order: Carnivora (a), Cetartiodactyla (b), Chiroptera (c), Primates (d), and Rodentia (e). Viral richness was calculated for each host order using two GAM models (both assuming maximum research effort). One class of models included variables related to historical mammal losses and the other class did not. The results from both models were visualized using IUCN range maps (2015.2), and for each mammalian order, the map generated without extinction variables was subtracted from the map that did, resulting in maps a – c.
